# Binding affinities of oxytocin, vasopressin, and Manning Compound at oxytocin and V1a receptors in Syrian hamster brains

**DOI:** 10.1101/2020.03.18.995894

**Authors:** Jack H Taylor, Katharine E McCann, Amy P Ross, H Elliott Albers

## Abstract

Oxytocin (OT) and arginine vasopressin (AVP), as well as synthetic ligands targeting their receptors (OTR, V1aR), are used in a wide variety of research contexts, but typically their pharmacological properties are determined in only a few species. Syrian hamsters (*Mesocricetus auratus*) have a long history of use as a behavioral and biomedical model for the study of oxytocin and vasopressin, and more recently, hamsters have been used to investigate behavioral consequences of OT-mediated activation of V1aRs. We sought to determine the binding affinities of OT, AVP, and the selective V1aR antagonist, Manning compound, in OTRs and V1aRs found in hamster brains. We performed saturation binding asays to determine the Kd values for the selective OTR and V1aR radioligands, [125I]OVTA and [125I]LVA in hamster brains. We then performed competition binding assays to determine Ki values for OT, AVP, and Manning compond at both the OTR and V1aR. We found that OT and AVP each had the highest affinity for their canonical receptors (OT-OTR Ki=4.28 nM; AVP-V1ar Ki=4.70 nM), and had the lowest affinity for their non-canonical ligands (OT-V1aR=495.2nM; AVP-OTR Ki=36.1 nM). Manning compound had the highest affinity for the V1aR (MC-V1aR Ki=6.87 nM; MC-OTR Ki=213.8 nM), but Manning compound was not as selective for the V1aR as has been reported in rat receptor. When comparing these data to previously published work, we found that the promiscuity of the V1aR in hamsters with respect to oxytocin and vasopressin binding is more similar to the promiscuity of the human V1aR than the rat V1aR receptor. Moreover, the selectivity of oxytocin at hamster receptors is more similar to the selectivity of oxytocin at human receptors than the selectivity of oxytocin at rat receptors. These data highlight the importance of determining the pharmacological properties of behaviorally relevant compounds in diverse models species.

---

The nonapeptide hormones oxytocin (OT) and arginine vasopressin (AVP) and their cognate receptors are widely used in a variety of therapeutic and research contexts, including the study of social behavior. Perhaps due to their roles in modulating a wide variety of social behaviors across mammalian species, OT and AVP have been studied in many species, each with its own behavioral, practical, and ethological considerations ^1,2^. Moreover, because OT and AVP can bind and activate both the OT receptor (OTR) and the AVP V1a receptor (V1aR) ^3^, there are dozens of pharmacological compounds that have been developed to selectively target each receptor ^4^. Despite the widespread study of OT and AVP, the binding affinities of OT and AVP with OTRs and V1aRs have been determined in only a handful of species, with the majority of data limited to human, mouse, and rat receptors ^4,5^ *c.f*. ^6,7^. Comparative studies, in which receptors from multiple species are probed at the same time using exactly the same methodology, show that there are differences in binding affinities (and subsequent activation potencies) even among closely related species ^6,7^. However, for many of the species that are currently being studied in the context of OT- and AVP-mediated social behavior, the binding affinities at the OTR and V1aR for these peptides, as well as for human-made pharmacological compounds, are not known.

The use of diverse animal models, each with its own unique constellation of social behavior, provides a comprehensive yet complex view of the roles of OT and AVP in modulating social behavior ^8^. A recent comparative analysis shows that, with respect to receptor distribution, OTR and V1aR are quite variable within *Rodentia* ^9^. Even among closely related species, there are differences in the OTR and V1aR amino acid sequences, which may translate to differences in ligand binding and signaling. Syrian hamsters are widely used as an animal model in social and biomedical research, and are preferred over other rodent models for many applications, including acute social stress ^10,11^, circadian rhythms ^12,13^, reproductive biology ^14^, sex differences in physiology and behavior ^15,16^, and infectious diseases ^17,18^. Hamsters are phylogenetically similar to rats and mice, but display distinct patterns of social behavior and ecology. Despite a long history of use in the context of AVP- and OT-mediated behavior ^19,20^, the binding affinities at the hamster OTR and V1aR for OT, AVP, and other compounds are not known. This is particularly notable given the potential for behavioral effects of OT acting at the V1aR, and AVP acting at the OTR, as it has been demonstrated that some V1aR-mediated behavior in this species may be via OT, and not AVP ^21^. Data regarding the binding affinities at the OTR and V1aR for these peptides, as well as specific agonists and antagonists, may hold explanatory power for *in vivo* behavioral pharmacology, particularly when AVP and OT are administered in regions in which OTR and V1aR distributions overlap, or when OT and AVP are compared to selective agonists and antagonists.

There are a variety of methods used to determine the binding affinity for ligands at g protein-coupled receptors (GPCRs), including radioligand binding assays (RBAs). All RBAs require some type of tissue that has been selectively enriched with the GPCR of interest. Isolated membrane and whole cultured cell preparations are popular methods, but an underused method of determining binding affinity is to carry out an RBA in intact, native tissue that is naturally enriched in the GPCR of interest. This method is simple, provides a cellular context that is as close as possible to *in vivo* conditions, and many laboratories that study OT- and AVP-mediated social behavior are already equipped to perform these RBAs. Here, we demonstrate the use of an RBA in intact brain sections to determine the binding affinities of the OTR and V1aR in Syrian hamsters. We predicted that if Syrian hamster OTRs and V1aRs function like rat and mouse receptors, then we should obtain binding affinities for the hamster receptors using various compounds that are on the same order of magnitude as have been reported for mouse and rat receptors. We tested the binding affinities for OT and AVP at their cognate receptors, as well as at each other’s receptors. We also tested the binding affinity of Manning Compound, a commonly used antagonist that is generally regarded as selective for V1aR in rat receptors, but not human or mouse ^4,5^.

## Methods

### Tissue preparation and quantification

Adult male hamsters were deeply anesthetized with isoflurane. Upon induction of complete analgesia (measured via a toe pinch), hamsters were decapitated and the brain was rapidly removed and snap frozen in isopentane chilled on dry ice. Brains were stored at −80° C until sectioning. Brains were hemisected, sectioned at 20 μm on a cryostat, thaw mounted onto Superfrost plus slides, and then returned to −80° C until the day of assay. Brains were cut into series of eight to nine, such that consecutive sections were mounted onto consecutive slides. Matching sets were generated by mounting contralateral sections on separate slides. On the day of assay, slides were allowed to come to room temperature and dry completely. Then, groups of four hemi-sections (n = 4 technical replicates) were roughly outlined with a wax pencil, and 250 μL of binding solution (radioligand plus 50 mM Trizma Base, 21mM MgCl, 1% (w/v) BSA, 0.5%(w/v) bacitracin, pH 7.4) was pipetted directly onto each set of four sections. Slides were left at room temperature for one hour, after which excess binding solution was removed, and the slides were washed via two five-minute washes and one 35-minute wash in buffer (50 mM Trizma Base, 21mM MgCl, pH 7.4). After washes, slides were allowed to dry completely, and then laid on film (Carestream BIOMAX MR) and given 2-3 days of exposure. All experiments were carried out on at least two different days using fresh aliquots of cold ligands. Images were captured using a camera mounted above a light box. Binding was quantified by comparing the exposure intensity of specifically bound nuclei to [14C] standards that were laid with each film, fitted to a Rodbard function. Radioactivity is expressed in the same units as the [14C] standards, DPM/0.4 mg. The nuclei that were quantified are shown in Figure 1. All animal procedures were carried out in accordance with the National Institutes of Health Guide for the Care and Use of Laboratory Animals and approved by the Georgia State University Institutional Animal Care and Use Committee.

**Figure 1.**
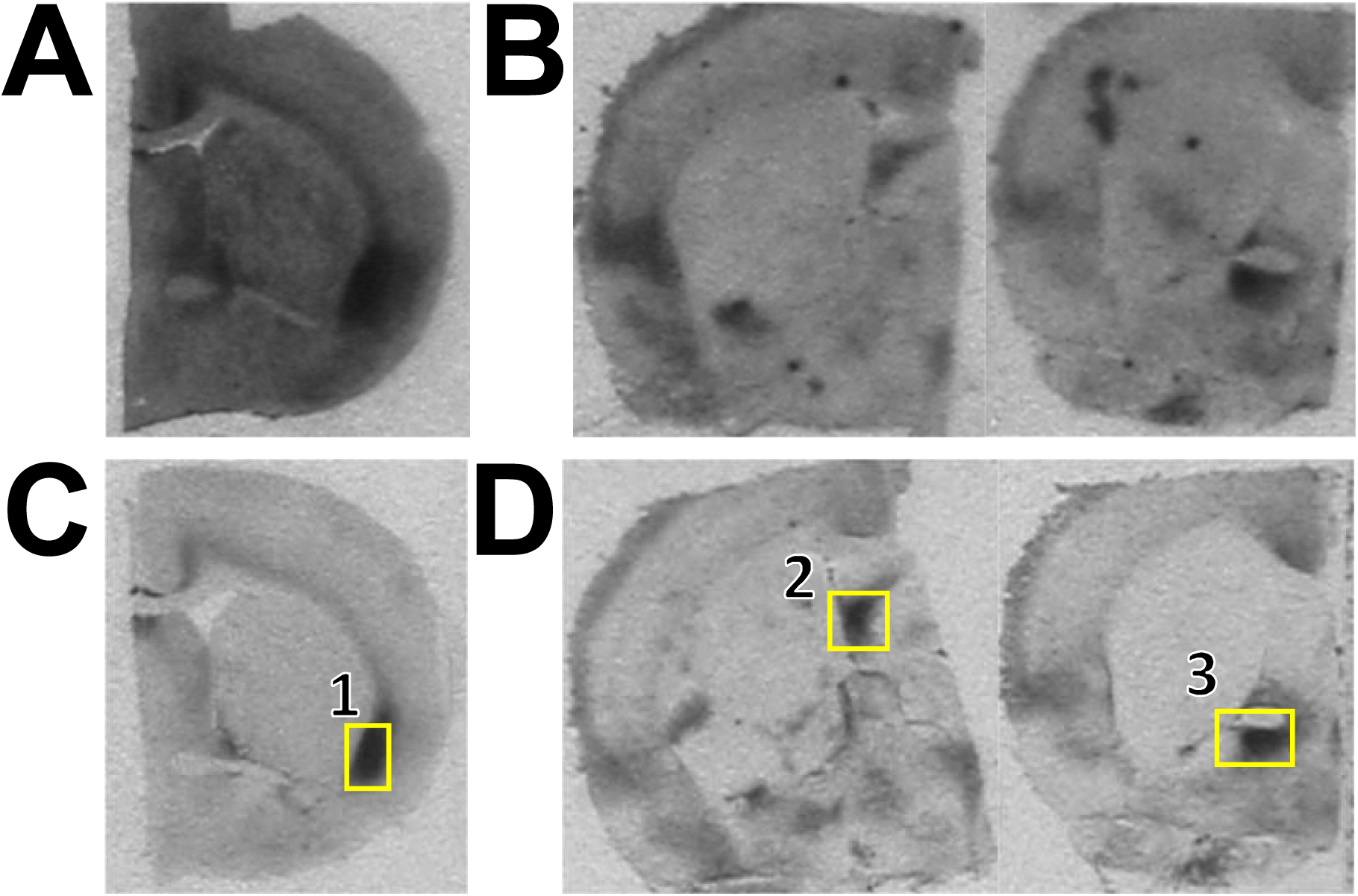
Representative photomicrographs of sections bound to A,C) [125I] OVTA or B,D) [125I] LVA. Each ligand produced distinct patterns of binding, even at A,B) 2000 pM concentrations. Regions quantified are shown in yellow boxes in C,D. 1. dorsal endopiriform nucleus, 2. lateral septum, 3. anteroventral bed nucleus of the stria terminalis

### Saturation Binding

Saturation binding experiments were carried out using eight to nine slide series. One hemisphere was used for total binding, while the contralateral section was used for non-specific binding. For OTRs, saturation binding occurred in the presence of doubling concentrations of [125I] OVTA (Perkin Elmer, NEX254010UC) ranging from 7.6 pM to 1000 pM. Non-specific binding was defined as 1 × 10^−5^ M OT. For V1aRs, saturation binding occurred in the presence of doubling concentrations of [125I] LVA (Perkin Elmer, NEX310050UC) ranging from 7.6 pM to 2000 pM. Non-specific binding was defined as 1 × 10^−5^ M AVP. Saturation curves were generated using curve fitting software (Graphpad Prism) and binding affinities (Kd) determined as the half maximal concentration. Reported Kd values and SEM were determined by averaging across biological replicates (n = 6-7). The potential for ligand depletion was assessed by pipetting 10 μL of each ligand concentration at the beginning and the end of the binding assay onto filter paper. These were then laid onto film with the tissue slides and compared for changes in film exposure.

### Competition Binding

Competition binding experiments were carried out using eight to nine slide series. For OTRs, competitive binding occurred using 200 pM [125I]OVTA in the presence or absence of competitor in concentrations ranging from 10^−11^-10^−4.5^ M. For V1aRs, competitive binding occurred using 400 pM [125I]LVA in the presence or absence of competitor in concentrations ranging from 10^−11^-10^−4.5^ M. Technical replicates (n = 4 hemisections per experiment) were averaged and treated as biological replicates (n = 4-9). Competition curves were generated using curve fitting software (Graphpad Prism) incorporating the measured Kd and concentration of the radioligand (i.e. a Cheng-Prusoff correction), and binding affinities (Ki) determined as the half maximal concentration. A Bonferroni-corrected cutoff (*p* = .05 ÷ 3 = .0167) was used to determine statistically significant differences in Ki values. The competitor ligands were OT (CYIQNCPLG-NH2 ; Bachem), AVP (CYFQNCPRG-NH2 ; Sigma) and Manning Compound (d(CH_2_)_5_[Tyr(Me)^2^]AVP ; Sigma)

## Results

### Saturation Binding

Saturation binding assays for [125I]OVTA and for [125I]LVA yielded Kd values that were similar to those reported in humans and rats (Table 1). The average Kd for [125I]OVTA in hamster tissue was 0.0945 (± .002) nM, with a Bmax of 2957 (± 55.03) DPM/ 0.04 mg. The average Kd for [125I]LVA in hamster tissue was 0.2619 (± 0.067) nM, with a Bmax of 5230 (± 1144) DPM/ 0.04 mg. Importantly, [125I]OVTA and [125I]LVA produced binding patterns that were visually distinct, even at the 2000 pM concentrations, indicating their selectivities as probes (Figure 1). Ligand depletion did not occur, as measured by 10 uL blots of radioligand buffer pipetted onto filter paper before and after binding had occurred. Representative saturation curves are shown in Figure 2.

**Table 1.**
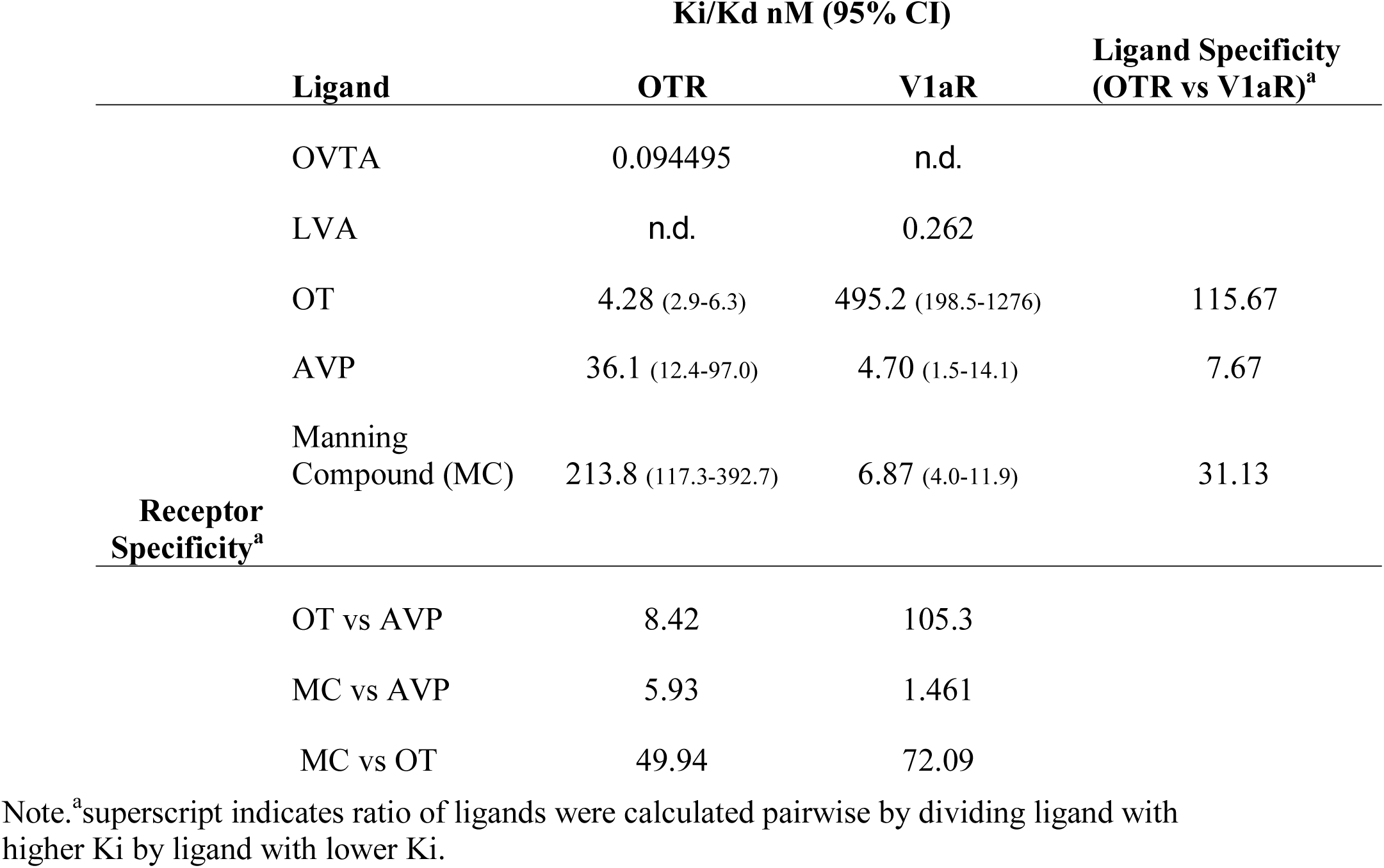
Binding affinities, ligand specificities, and receptor specificities.

**Figure 2.**
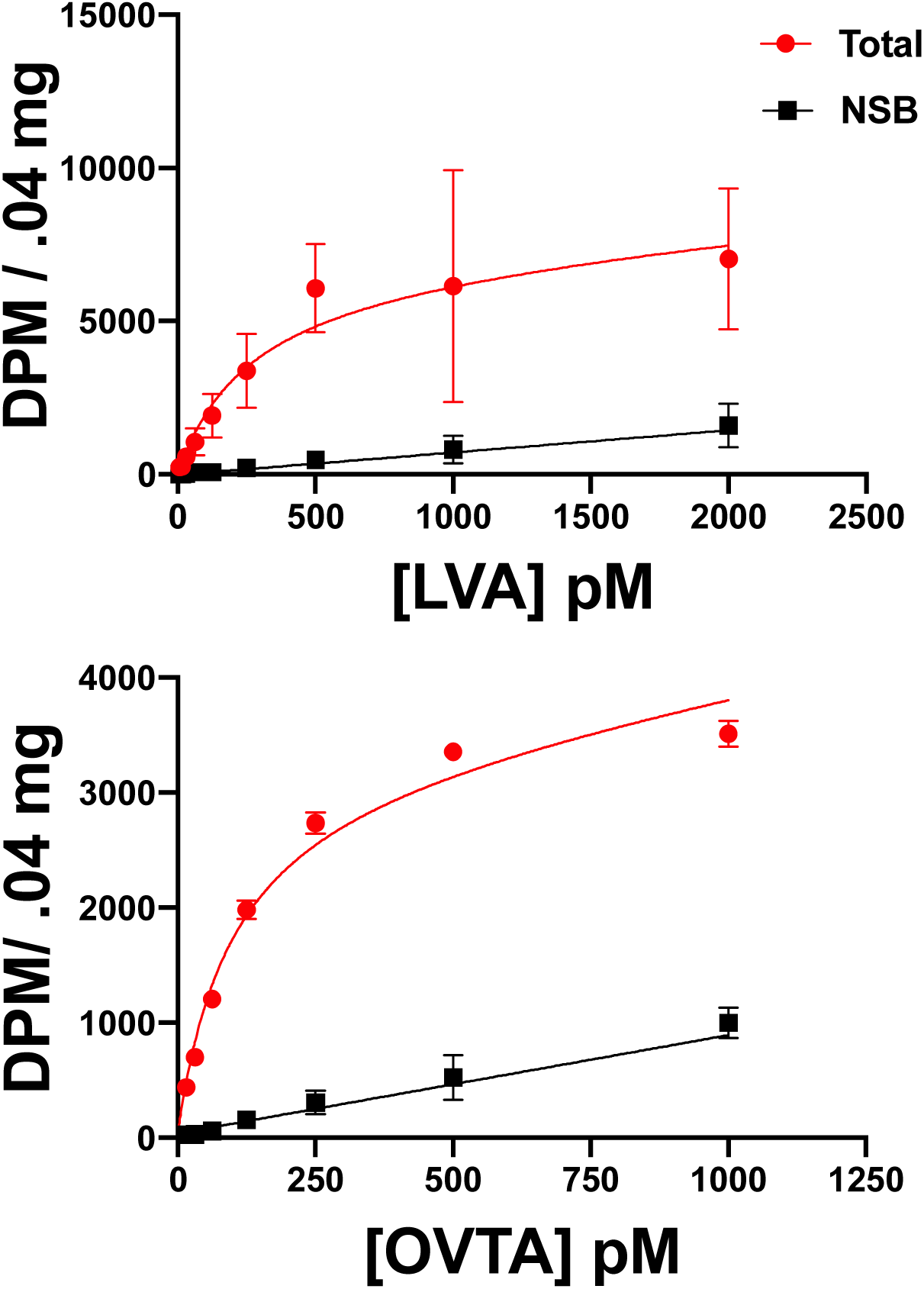
Representative saturation curves for [125I] LVA (top) and [125I] OVTA (bottom) in hamster brain slices. Total = Total binding, NSB = Non Specific Binding

### Competitive Binding

Competitive binding assays yielded Ki values for each ligand that were within the expected ranges, based on previous reports in other species (Table 1, Figure 3). At the hamster OTR, OT had an average Ki of 4.28 nM, AVP had an average Ki of 36.06 nM, and Manning Compound had a Ki of 213.8 nM. The binding affinity of OT at the OTR was significantly lower than that of AVP (F_1,140_ = 17.55, *p* < .0001) and Manning compound (F_1,119_ = 61.94, *p* < .0001). The binding affinity of AVP at the OTR was significantly lower than that of Manning compound (F_1,111_ = 17.55, *p* = .0148). At the V1aR, OT had a Ki of 495.2 nM, AVP had a Ki of 4.70 nM, and Manning Compound had an average Ki of 6.87 nM. The binding affinity of AVP at the V1aR was significantly lower than that of OT (F_1,88_ = 13.37, *p* = .0004), but was not significantly different than that of Manning compound (F_1,82_ = 224, *p* < .638). The binding affinity of Manning compound at the V1aR was significantly lower than that of OT (F_1,60_ = 55.39, *p* < .0001). Importantly, these data show that in hamsters, OT and AVP bind with the highest affinity (i.e. lowest Ki) to their cognate receptors.

**Figure 3.**
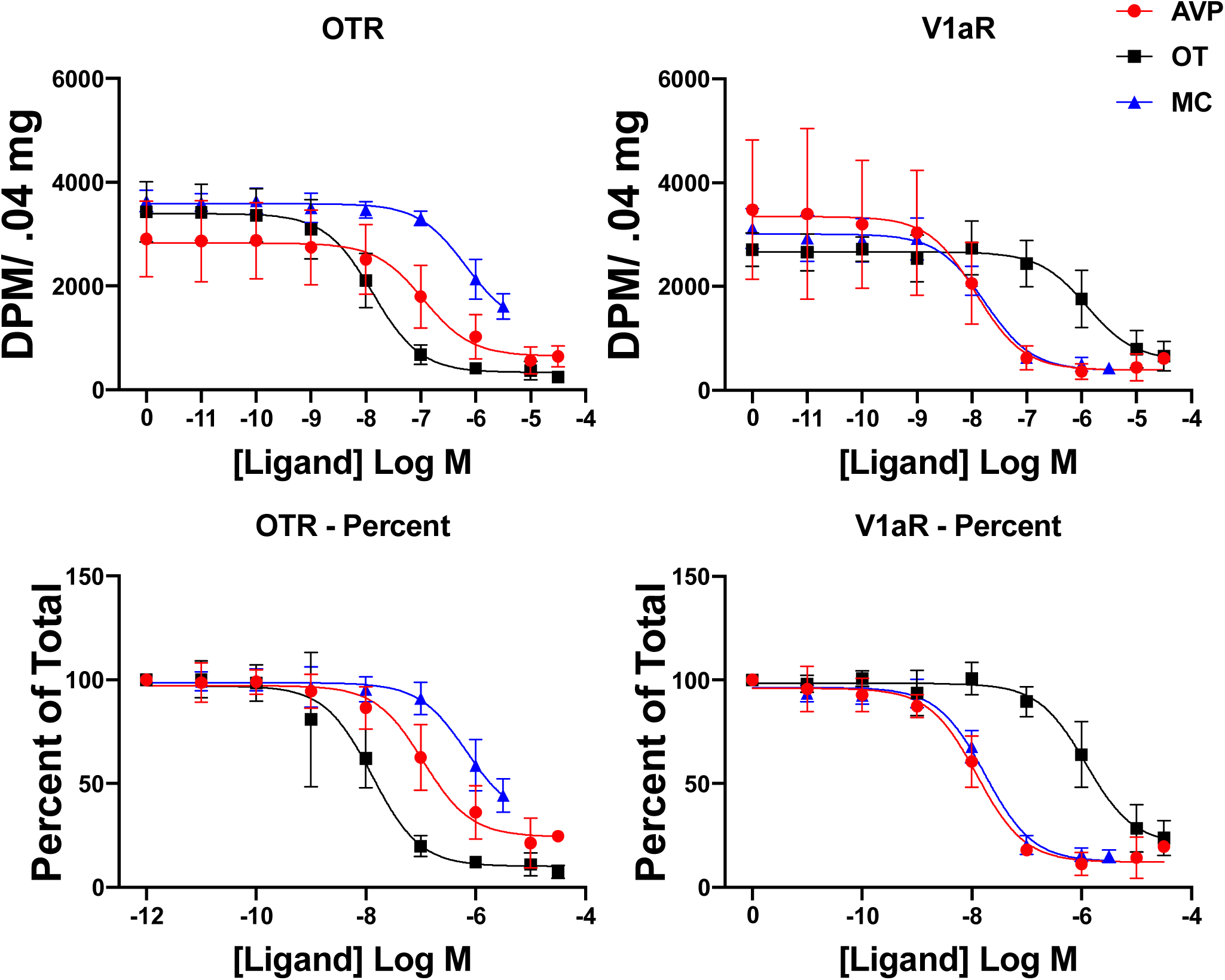
Competition curves for oxytocin (OT) vasopressin (AVP) and Manning compound (MC) at the hamster OT receptor (left) and vasopressin V1a receptor (right).

## Discussion

The data presented here indicate that many properties of V1aRs and OTRs in Syrian hamsters more closely resemble those of human receptors than those of more closely related rodent species (i.e. rats and mice), and demonstrate the usefulness of hamsters and less commonly used animal models for pharmacological experiments. Translational relevance is crucial for preclinical research using animal models. While mouse and rat models of human behavior and disease are beneficial because of the plethora of genetic tools available, these models often fall short of providing ethologically- or translationally-relevant readouts. Thus, the use of diverse model systems is necessary for providing a better understanding of the molecular mechanisms underlying behavior and disease. For examples, hamsters are a preferred model for investigating reproductive biology ^14^, tumor growth ^22,23^, and more recently in studying properties of infectious diseases, including Ebola, Zika, and influenza viruses ^17,18,24,25^. Furthermore, the human stress response system is more similar to the hamster system than to that of other rodent models and both male and female hamsters readily exhibit aggressive behavior, making hamsters an excellent model to investigate behavioral, physiological, and molecular responses to social stress in both sexes ^10,11,16^. The data presented here indicate that many properties of V1aR and OTR in Syrian hamsters more closely resemble those of human receptors than those of more closely related rodent species, suggesting that further work in hamsters may help to reveal more translationally-relevant details on how these ligands interact with their receptors to facilitate behavior.

While it is not prudent to directly compare the binding affinities obtained here to the binding affinities obtained through other methods, it is valuable to compare the *relationships* between binding affinities of different ligands at different receptors. For example, our results indicate that the degree of promiscuity of the hamster V1aR, but not the hamster OTR, is more similar to the promiscuity of human receptors than to rat receptors. We found that the affinity of OT at the OTR was ∼8.4 fold better than the affinity of AVP at the OTR. In rats, this same relationship is only ∼1.7 fold, while in humans, it is ∼2.1 fold different ^4^. Likewise, the affinity of AVP at the hamster V1aR was ∼105 fold better than the affinity of OT at the V1aR. In rats, this relationship is only ∼27.3 fold, while in humans and other primates it is >100 fold ^4,6,7^. Interestingly, when focusing on the selectivity of the ligands rather than the selectivity of the receptors, we found that the selectivity of OT, but not AVP in hamsters more closely resembles that in humans than rats. OT is ∼116 fold more selective for the hamster OTR than for the V1aR. In rats, this relationship is ∼71 fold, while in humans it is ∼150 fold different ^4^. Likewise, AVP is ∼7.7 fold more selective for the hamster V1aR than for the OTR. In rats, this relationship is ∼2.167 fold, while in humans it is only ∼1.54 fold. Thus, with respect to the promiscuity of the *receptors*, the OT/AVP relationship in the hamster V1aR is more similar to the same relationship in humans than in rats; however, with respect to the *ligands*, the selectivity of OT in hamsters is more similar to the same relationship in humans than in rats. Furthermore, Manning compound is ∼31 fold more selective for the hamster V1aR than for the OTR, which is not considered “selective” by the definition of a 100 fold difference in selectivity used by ^4^, a criteria which Manning compound meets in rat receptors. However, this compound is more selective for the V1aR in hamsters than it is in humans, and slightly more selective than it is in mice ^5^. Taken together, these findings highlight the need to carefully evaluate a ligand’s pharmacological properties when applying it to new model species, whether that be through an RBA or through other bio- or behavioral assays.

In this paper we showed that the binding affinities for commonly used pharmacological compounds can be quickly and easily measured in a species for which enriched tissue is available. Many labs that regularly utilize *in vivo* behavioral pharmacology already have the capacity to perform similar assays using their species and ligands of choice. As the number of model species for various behavioral inquiries grows, it is important to consider the possibility that ligands which are validated for use in mice and rats may not act on other species’ receptors in the same way, even in species that are phylogenetically similar. This method also provides a cellular context that is as close as possible to the binding conditions in living, native tissue. This is an important factor to consider, because there is the possibility that receptor heterodimers influenced the binding affinities of each ligand at the OTR or V1aR. Both the OTR and V1aR form heterodimers with each other, as well as with the vasopressin V2 receptor, but these heterodimers exhibit only small differences in binding affinity for AVP ^26^. When all of these considerations are taken together, the potential for a myriad of complex influences makes this method an attractive alternative and adjunct to other RBA methods for studying ligand-receptor dynamics under conditions that are as close as possible to those *in vivo*.

Though these data provide new information on the pharmacological characteristics of commonly used ligands in a widely used model species, they do not provide any information on the ability of OT, AVP, or Manning Compound to activate one or more signaling cascades. Binding affinity and selectivity are the critical factors in the pharmacological properties of a true antagonist, but for agonists and inverse agonists, binding affinity and receptor activation can often tell quite different stories. For example, the binding affinity for AVP at the OTR for human and macaque receptors is similar, but AVP exhibits nearly 10 fold different potency for calcium mobilization between these two species ^7^. Likewise, the affinity of OT at the human V1aR is ∼15 fold better than the affinity of OT at the marmoset V1aR, but OT acts as a partial agonist/ partial antagonist for calcium mobilization at the human V1aR, and a superagonist at the marmoset V1aR ^6^. Thus, just as it is important to evaluate how well ligands bind to the receptors of various animal models, it is also important to evaluate the cellular and molecular consequences of ligand binding.

We have shown here that the relationship in hamsters between OT, AVP, and their receptors in some ways more closely resembles those same relationships in humans than in rats, highlighting important differences in binding patterns of closely related species. The determination of the pharmacological properties of both native ligands and those used as research tools is an important component to understanding how these ligands affect behavior. As more diverse animal models continue to be developed, it will be critical to re-evaluate the compounds that are used, as well as highlight the differences and similarities between species and how these variations in pharmacological properties correlate to social behavior. In particular, comparative approaches to pharmacological experiments have the potential to provide a wealth of information that can help when translating to human research.

## Acknowledgements

The authors would like to thank Alisa Norvelle, M.S. for assistance performing the experimental procedures. This research was supported by NIH grant MH110212 to HEA.

## Data Availability Statement

Data are available upon reasonable request to the corresponding or senior authors.

## References

1. Dumais, K. M. & Veenema, A. H. Vasopressin and oxytocin receptor systems in the brain: Sex differences and sex-specific regulation of social behavior. Frontiers in Neuroendocrinology 40, 1–23 (2016).

2. Jurek, B. & Neumann, I. D. The Oxytocin Receptor: From Intracellular Signaling to Behavior. Physiological Reviews 98, 1805–1908 (2018).

3. Song, Z. & Albers, H. E. Cross-talk among oxytocin and arginine-vasopressin receptors: Relevance for basic and clinical studies of the brain and periphery. Front Neuroendocrinol 51, 14–24 (2018).

4. Manning, M. et al. Oxytocin and Vasopressin Agonists and Antagonists as Research Tools and Potential Therapeutics. Journal of Neuroendocrinology 24, 609–628 (2012).

5. Busnelli, M., Bulgheroni, E., Manning, M., Kleinau, G. & Chini, B. Selective and Potent Agonists and Antagonists for Investigating the Role of Mouse Oxytocin Receptors. J Pharmacol Exp Ther 346, 318–327 (2013).

6. Mustoe, A., Schulte, N. A., Taylor, J. H., French, J. A. & Toews, M. L. Leu 8 and Pro 8 oxytocin agonism differs across human, macaque, and marmoset vasopressin 1a receptors. Sci Rep 9, 1–10 (2019).

7. Taylor, J. H., Schulte, N. A., French, J. A. & Toews, M. L. Binding Characteristics of Two Oxytocin Variants and Vasopressin at Oxytocin Receptors from Four Primate Species with Different Social Behavior Patterns. J Pharmacol Exp Ther 367, 101–107 (2018).

8. Albers, H. E. Species, sex and individual differences in the vasotocin/vasopressin system: Relationship to neurochemical signaling in the social behavior neural network. Frontiers in Neuroendocrinology 36, 49–71 (2015).

9. Freeman, A. R., Aulino, E. A., Caldwell, H. K. & Ophir, A. G. Comparison of the distribution of oxytocin and vasopressin 1a receptors in rodents reveals conserved and derived patterns of nonapeptide evolution. Journal of Neuroendocrinology n/a, e12828 (2020).

10. Huhman, K. L. et al. Conditioned defeat in male and female syrian hamsters. Hormones and Behavior 44, 293–299 (2003).

11. Potegal, M., Huhman, K., Moore, T. & Meyerhoff, J. Conditioned defeat in the Syrian golden hamster (Mesocricetus auratus). Behavioral and Neural Biology 60, 93–102 (1993).

12. Albers, H. E. Response of hamster circadian system to transitions between light and darkness. American Journal of Physiology-Regulatory, Integrative and Comparative Physiology 250, R708–R711 (1986).

13. Huhman, K. L. & Albers, H. E. Neuropeptide Y microinjected into the suprachiasmatic region phase shifts circadian rhythms in constant darkness. Peptides 15, 1475–1478 (1994).

14. Hirose, M. & Ogura, A. The golden (Syrian) hamster as a model for the study of reproductive biology: Past, present, and future. Reproductive Medicine and Biology 18, 34–39 (2019).

15. Hennessey, A. C., Huhman, K. L. & Albers, H. E. Vasopressin and sex differences in hamster flank marking. Physiology & Behavior 55, 905–911 (1994).

16. McCann, K. E., Sinkiewicz, D. M., Rosenhauer, A. M., Beach, L. Q. & Huhman, K. L. Transcriptomic Analysis Reveals Sex-Dependent Expression Patterns in the Basolateral Amygdala of Dominant and Subordinate Animals After Acute Social Conflict. Mol. Neurobiol. 56, 3768–3779 (2019).

17. Iwatsuki-Horimoto, K. et al. Syrian Hamster as an Animal Model for the Study of Human Influenza Virus Infection. Journal of Virology 92, (2018).

18. Miao, J., Chard, L. S., Wang, Z. & Wang, Y. Syrian Hamster as an Animal Model for the Study on Infectious Diseases. Front. Immunol. 10, (2019).

19. Albers, H. E. & Ferris, C. F. Behavioral effects of vasopressin and oxytocin within the medial preoptic area of the golden hamster. Regulatory Peptides 12, 257–260 (1985).

20. Ferris, C. F., Albers, H. E., Wesolowski, S. M., Goldman, B. D. & Luman, S. E. Vasopressin injected into the hypothalamus triggers a stereotypic behavior in golden hamsters. Science 224, 521–523 (1984).

21. Song, Z. et al. Oxytocin induces social communication by activating arginine-vasopressin V1a receptors and not oxytocin receptors. Psychoneuroendocrinology 50, 14–19 (2014).

22. Gimenez-Conti, I. B. & Slaga, T. J. The hamster cheek pouch carcinogenesis model. J. Cell. Biochem. Suppl. 17F, 83–90 (1993).

23. Thomas, M. A. et al. Syrian hamster as a permissive immunocompetent animal model for the study of oncolytic adenovirus vectors. Cancer Res. 66, 1270–1276 (2006).

24. Miller, L. J. et al. Zika Virus Infection in Syrian Golden Hamsters and Strain 13 Guinea Pigs. Am. J. Trop. Med. Hyg. 98, 864–867 (2018).

25. Wahl-Jensen, V. et al. Use of the Syrian hamster as a new model of ebola virus disease and other viral hemorrhagic fevers. Viruses 4, 3754–3784 (2012).

26. Terrillon, S. et al. Oxytocin and Vasopressin V1a and V2 Receptors Form Constitutive Homo- and Heterodimers during Biosynthesis. Molecular Endocrinology 17, 677–691 (2003).

